# Polygenic scores for major depressive disorder and depressive symptoms predict response to lithium in patients with bipolar disorder

**DOI:** 10.1101/449363

**Authors:** Azmeraw T. Amare, Klaus Oliver Schubert, Liping Hou, Scott R. Clark, Sergi Papiol, Micah Cearns, Urs Heilbronner, Franziska Degenhardt, Fasil Tekola-Ayele, Yi-Hsiang Hsu, Tatyana Shekhtman, Mazda Adli, Nirmala Akula, Kazufumi Akiyama, Raffaella Ardau, Bárbara Arias, Jean-Michel Aubry, Lena Backlund, Abesh Kumar Bhattacharjee, Frank Bellivier, Antonio Benabarre, Susanne Bengesser, Joanna M. Biernacka, Armin Birner, Clara Brichant-Petitjean, Pablo Cervantes, Hsi-Chung Chen, Caterina Chillotti, Sven Cichon, Cristiana Cruceanu, Piotr M. Czerski, Nina Dalkner, Alexandre Dayer, Maria Del Zompo, J. Raymond DePaulo, Bruno Étain, Peter Falkai, Andreas J. Forstner, Louise Frisen, Mark A. Frye, Janice M. Fullerton, Sébastien Gard, Julie S. Garnham, Fernando S. Goes, Maria Grigoroiu-Serbanescu, Paul Grof, Ryota Hashimoto, Joanna Hauser, Stefan Herms, Per Hoffmann, Andrea Hofmann, Stephane Jamain, Esther Jiménez, Jean-Pierre Kahn, Layla Kassem, Po-Hsiu Kuo, Tadafumi Kato, John Kelsoe, Sarah Kittel-Schneider, Sebastian Kliwicki, Barbara König, Ichiro Kusumi, Gonzalo Laje, Mikael Landén, Catharina Lavebratt, Marion Leboyer, Susan G. Leckband, Alfonso Tortorella, Mirko Manchia, Lina Martinsson, Michael J. McCarthy, Susan McElroy, Francesc Colom, Marina Mitjans, Francis M. Mondimore, Palmiero Monteleone, Caroline M. Nievergelt, Markus M. Nöthen, Tomas Novák, Claire O’Donovan, Norio Ozaki, Urban Ösby, Andrea Pfennig, James B. Potash, Andreas Reif, Major Depressive Disorder Working Group of the Psychiatric Genomics Consortium, Eva Reininghaus, Guy A. Rouleau, Janusz K. Rybakowski, Martin Schalling, Peter R. Schofield, Barbara W. Schweizer, Giovanni Severino, Paul D. Shilling, Katzutaka Shimoda, Christian Simhandl, Claire M. Slaney, Alessio Squassina, Thomas Stamm, Pavla Stopkova, Mario Maj, Gustavo Turecki, Eduard Vieta, Julia Veeh, Stephanie H. Witt, Adam Wright, Peter P. Zandi, Philip B. Mitchell, Michael Bauer, Martin Alda, Marcella Rietschel, Francis J. McMahon, Thomas G. Schulze, Bernhard T. Baune

## Abstract

**Background:** Lithium is a first-line medication for bipolar disorder (BD), but only ~30% of patients respond optimally to the drug. Since genetic factors are known to mediate lithium treatment response, we hypothesized whether polygenic susceptibility to the spectrum of depression traits is associated with treatment outcomes in patients with BD. In addition, we explored the potential molecular underpinnings of this relationship.

**Methods:** Weighted polygenic scores (PGSs) were computed for major depressive disorder (MDD) and depressive symptoms (DS) in BD patients from the Consortium on Lithium Genetics (ConLi^+^Gen; n=2,586) who received lithium treatment. Lithium treatment outcome was assessed using the ALDA scale. Summary statistics from genome-wide association studies (GWAS) in MDD (130,664 cases and 330,470 controls) and DS (n=161,460) were used for PGS weighting. Associations between PGSs of depression traits and lithium treatment response were assessed by binary logistic regression. We also performed a cross-trait meta-GWAS, followed by Ingenuity^®^ Pathway Analysis.

**Outcomes:** BD patients with a low polygenic load for depressive traits were more likely to respond well to lithium, compared to patients with high polygenic load (MDD: OR =1.64 [95%CI: 1.26-2.15], lowest vs highest PGS quartiles; DS: OR=1.53 [95%CI: 1.18-2.00]). Associations were significant for type 1, but not type 2 BD. Cross-trait GWAS and functional characterization implicated voltage-gated potassium channels, insulin-related pathways, mitogen-activated protein-kinase (MAPK) signaling, and miRNA expression.

**Interpretation:** Genetic loading to depression traits in BD patients lower their odds of responding optimally to lithium. Our findings support the emerging concept of a lithium-responsive biotype in BD.

**Funding:** See attached details

## Introduction

Bipolar disorder (BD) is a chronic and severe psychiatric illness characterized by episodic, abnormal manic and depressive mood states. An estimated 48.8 million people are affected by BD globally^1^. The disorder accounts for 9.9 million years of life lived with disability worldwide^1^, and substantially increases all-cause mortality and risk of suicide^2^. Amongst available treatments, lithium is regarded as a gold standard by several clinical guidelines^3,4^. Lithium uniquely protects against both manic and depressive illness phases, has demonstrated protective effects against suicide^5-7^, and is particularly effective in preventing rehospitalisation^8^. However, not all patients with BD fully benefit from lithium, and only about 30% show full response to the drug^5-7^. In current psychiatric practice, no biological or clinical markers exist that could reliably predict responsiveness to lithium^9^, and prescribing cannot be targeted to patients who benefit most while avoiding side effects and sub-optimal treatment for poor responders^10,11,12,13^.

In order to develop objective response markers and to move towards personalized prescribing of lithium for BD patients, a better understanding of the biological mechanisms underlying lithium response is urgently required. Recent genome-wide association studies (GWAS) carried out by our International Consortium on Lithium Genetics (ConLi^+^Gen)^5^ and others^14,15^ have indicated that genetic variation could be an important mediator of response to long-term lithium treatment response in BD patients. Additionally, we have recently demonstrated that high genetic loading for schizophrenia (SCZ) risk variants in people with BD decreases the likelihood of favorable response to lithium^16^, suggesting that polygenic score (PGS) analysis of mental and physical traits could yield important information on the genetic architecture of BD phenotypes^17,18,19^. In the current study, we address the question of whether genetic loading for major depressive disorder (MDD) and depressive symptoms (DS) contribute to treatment outcomes in BD.

BD and MDD show 47% genetic overlap^20-22^, and shared risk genes and biological pathways have been described^22,23^. Lithium can be effective as an augmentation strategy in MDD patients who have experienced an insufficient response to first-line antidepressants^24,25^ and is protective against further MDD episodes after symptom remission has been achieved^20,26^. Moreover, a large observational study based on the Finnish registry showed that lithium is the most effective agent preventing rehospitalization in MDD^26^.

On the other hand, in BD, lithium is more effective in preventing manic than depressive episodes^27,28^, leading to the notion that better lithium responders might be more likely to experience manic predominant polarity, as opposed to depressive predominant polarity^29^. In support of this view, one study found that excellent lithium responders were characterized by a manic but not depressive polarity of the index episode ^30^. Another study described an episodic illness pattern of ‘mania-depression-interval’ as a predictor for good response, whereas a ’depression-mania-interval’ predicted poorer outcomes^31^. Inter-episode residual mood symptoms, as opposed to full remission^6,7,32^, a rapid cycling pattern^31,32^, and a history of mixed episodes^33,34^ have also been described as predictors of poor response.

On the background of these complex interactions between BD, MDD, and lithium treatment, we asked whether BD patients with a high genetic susceptibility for depression (MDD and DS), expressed by their PGS for these traits, would respond better or worse to lithium than BD patients with a low genetic loading^35,36^. To explore potential genetic and molecular drivers of any detected polygenic association, we carried out a cross-trait GWAS meta-analysis, combining summary statistics from the largest available GWASs for MDD^35^ and DS^36^ with GWASs for response to lithium treatment in patients with BD^5^. Overlapping SNPs that met genome-wide significance in the meta-GWAS were subsequently analyzed for biological context using the Ingenuity^®^ Pathway Analysis platform (IPA^®^).

## Methods and Materials

### Discovery GWAS summary data sets

The polygenic score and cross-trait meta-analysis for this study were based on genetic data from the International Consortium on Lithium Genetics (ConLi^+^Gen)^5^, and the summary statistics of three largest GWASs available for MDD^35^, DS^36^ and treatment response to lithium in patients with BD^5^.

#### Major depressive disorder

The most recent GWAS meta-analysis of 9.6 million SNPs (Psychiatric Genomics Consortium-PGC; http://www.med.unc.edu/pgc/), obtained from 7 cohorts (deCODE, Generation Scotland, GERA, iPSYCH, UK Biobank, CONVERGE and 23andMe) containing 130,664 MDD cases and 330,470 healthy controls, identified 44 independent loci that reached the criteria for statistical significance. Details on this study are available elsewhere^35^.

#### Depressive symptoms

The GWAS on DS (N = 161,460) used data from the PGC, the UK Biobank (UKB), the Resource for Genetic Epidemiology Research on Aging (GERA) Cohort and the Social Science Genetic Association Consortium (SSGAC) https://www.thessgac.org/. The summary statistics were made publically available for scientific usage^36^.

#### Lithium treatment response in BD

The summary GWAS on lithium treatment response was produced through a combined analysis of 2,563 patients collected by 22 participating sites from the International Consortium on Lithium Genetics (ConLi^+^Gen) http://www.conligen.org/. In our analysis, we used the data analyzed on the categorical scale for lithium response^5^.

### Target Study Sample

For the PGS analysis, clinical data on lithium treatment response and genetic information were obtained <u>from</u> the International Consortium on Lithium Genetics (ConLi^+^Gen; www.ConLiGen.org) for n=2,586 patients (including 23 patients in the replication sample)^3,5,16^. A series of quality control procedures were implemented on the genotype data before and after imputation as described below.

### Target outcome

Lithium treatment response was assessed using the validated “Retrospective Criteria of Long-Term Treatment Response in Research Subjects with Bipolar Disorder” scale, also known as the ALDA scale ^7,37,38^. This scale quantifies symptom improvement over the course of treatment (A score, range 0–10), which is then weighted against five criteria (B score) that assess confounding factors^5^. Patients with a total score of 7 or higher were categorized as “good responders”, and the remainder were categorized as poor responders^5,38^. In addition to the ALDA scale scores, information on covariates such as age and gender was collected, as described in detail elsewhere^5^.

### Genotyping and quality control

The genome-wide genotypes, as well as clinical and demographic data, were collected by 22 participating sites. Quality control (QC) procedures were implemented on the genotype data using PLINK, version 1.09 prior to imputation^39^. Samples with low genotype rates <95%, sex inconsistencies (based on X-chromosome heterozygosity), and one of a pair of genetically related individuals were excluded. SNPs were excluded based on the following criteria: a poor genotyping rate (<95%), strand ambiguity (A/T and C/G SNPs), a low minor allele frequency (MAF<1%), or those deviated from genotype frequency expectations under the Hardy-Weinberg Equilibrium (p<10^−6^).

### Imputation

The genotype data passing QC were imputed on the Michigan server^40^ (https://imputationserver.sph.umich.edu) separately for each genotype platform using reference data from the 1000 Genomes Project Phase 3 (Version 5). The European reference panel was used for all the samples except for those from Japan and Taiwan, for which an East Asian reference population data was used. After excluding low-frequency SNPs (MAF<10%); low-quality variants (imputation INFO < 0.9); and indels, the imputed dosages were converted to best guess genotypes. The subsequent polygenic analyses were performed using these best guess genotypes.

## Statistical Analyses

### Polygenic score (PGS) association analysis

PGSs were calculated using the approach previously described by the International Schizophrenia Consortium^41^. Prior to PGS computation, independent SNPs were identified through a clumping procedure implemented in PLINK software, version 1.09 run on Linux^39^. Quality-controlled SNPs were clumped for linkage disequilibrium based on GWAS association p-value informed clumping at r^2^ = 0.1 within a 250- kilobase window to create a SNP-set in linkage equilibrium (*plink-clump-p1 1-clump-p2 1-clump-r2 0.1-clump-kb 250*). Polygenic risk scores were calculated for MDD and DS in the ConLi^+^Gen sample at ten GWAS association p-value thresholds (<1 × 10^−4^, <1 × 10^−3^, <0.01, <0.05, <0.1, <0.2, <0.3, <0.4, <0.5, <1).

### Cross-trait meta-analysis of genome-wide association studies

Having identified a significant polygenic association that indicated the presence of genetic overlap, we conducted cross-trait meta-analyses of GWASs to identify genetic polymorphisms that were likely to increase the susceptibility to both MDD and DS as well as influence lithium treatment response in patients with BD. The cross-trait meta-analyses were performed by combining the summary statistics for GWAS on lithium response ^5^ and GWAS on MDD^35^ and DS^36^. We applied the O‘Brien’s (OB) method and the direct Linear Combination of dependent test statistics (dLC) approach^42,43^, which are implemented in the C^++^ eLX package (further details in supplementary methods).

### Ingenuity^®^ Pathway Analysis (IPA^®^)

To characterize the biological context of the discovered SNPs from the cross-trait meta-analyses, we implemented a functional analysis using QIAGEN’s Ingenuity^®^ Pathway Analysis (IPA***^®^***, QIAGEN Redwood City, CA, USA, www.qiagen.com/ingenuity). For details see supplementary methods.

## Results

### Sample characteristics and lithium treatment response rate

After QC, 2,586 patients (3,193 before QC) remained for analysis. While 2,366 were of European ancestry, the remaining were of Asian ancestry. In all, 704 (27.2%) responded to lithium treatment (ALDA score ≥7). Detailed sample and demographics details have been described previously^16^.

### MDD and DS PGS are associated with lithium treatment response in BD

Associations between the PGSs for MDD and DS with lithium treatment response were found at various p-value thresholds. The strongest association were found for MDD (p= 0.0003) at P_T_ <5 × 10^−2^, R^2^ = 0.7% and for DS (p= 0.0003) at P_T_ <1 × 10^−2^, R^2^ = 0.7%) (Figure 1).

**Figure 1.**
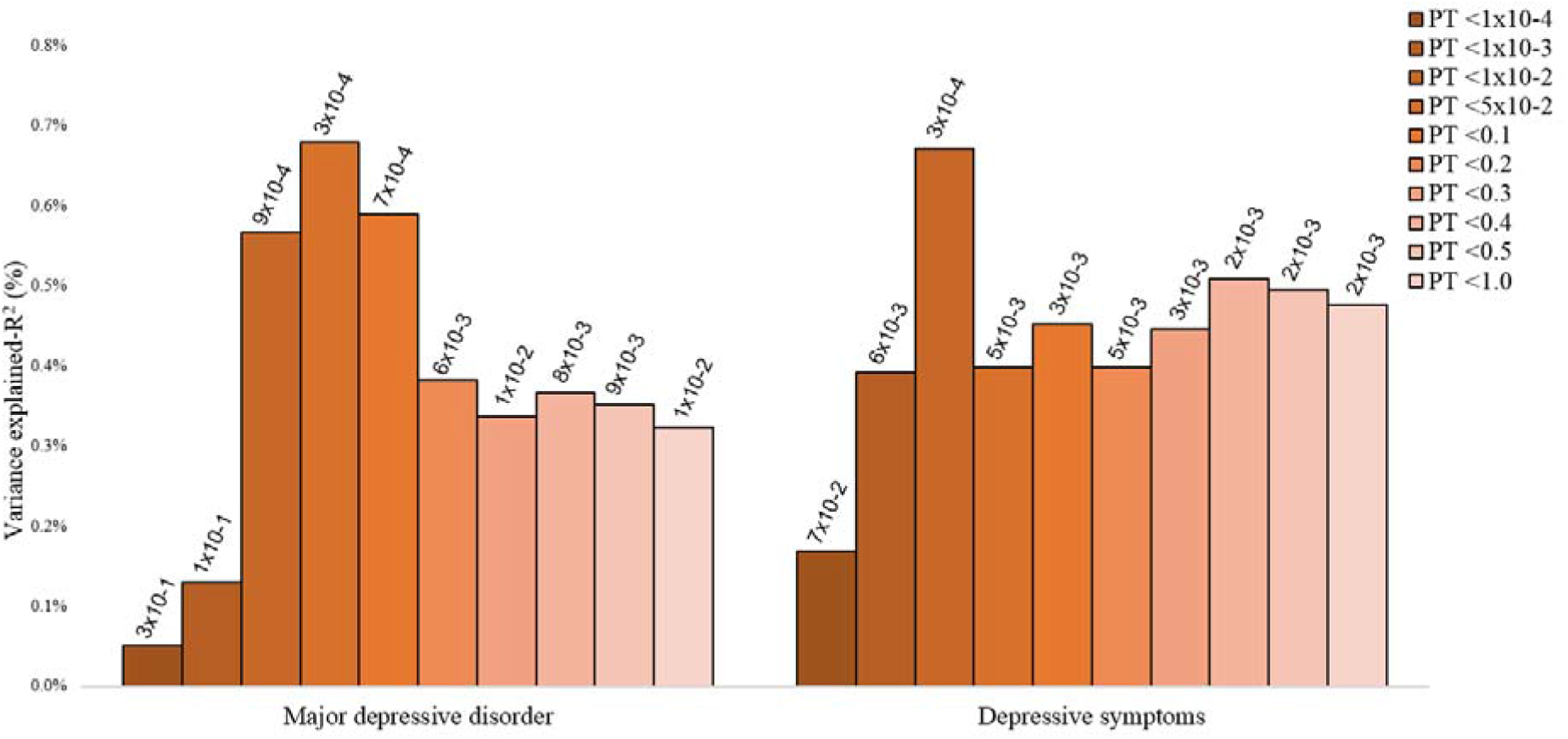
The association of PGS for depression traits (MDD and DS) and lithium treatment response at different GWAS p-value thresholds. The y-axis refers to the percentage of variance in treatment response to lithium accounted for by the PGSs for depressive traits at particular P-value thresholds. On the x-axis, are the GWAS P-value thresholds used to select single-nucleotide polymorphisms for the PGSs. On the top of each bar are the p-values for the association between the PGSs for depressive traits and lithium treatment response.

### High genetic loadings for MDD and DS are associated with poorer response to lithium in BD

We divided the study population into quartiles, according to their polygenic loading for MDD and DS, respectively. As shown in Figure 2 and Table 1, BD patients who carry a lower polygenic load (1^st^ quartile) for MDD or DS have higher odds of favorable lithium treatment response, compared to patients carrying a high polygenic load (4^th^ quartile). The odds ratio (OR) of favorable response for patients in the 1^st^ quartile compared with those in the 4^th^ quartile was 1.64 [95%CI: 1.26-2.15] for MDD PGS, and 1.53 [95%CI: 1.18-2.00]) for DS PGS (Table 1 & Figure 2).

**Table 1.** Odds ratios of favorable lithium treatment response in patients with BD, comparing the response status of patients in the low PGS quartile for MDD and DS with patients with the highest polygenic load for MDD/DS (4^th^ quartile).

| Categories | Patients with BD (n=2,586) |  |  |
| --- | --- | --- | --- |
|  | Responders/<br>Non-responders | unadjusted OR<br>(95% CI) | ¥Adjusted OR<br>(95% CI) |
| MDD |  |  |  |
| **1<sup>st</sup> lowest score** | 197/449 | 1.56 (1.21-2.00) | 1.64 (1.26-2.15) |
| **2<sup>nd</sup>** | 202/444 | 1.62 (1.26-2.07) | 1.60 (1.22-2.08) |
| **3<sup>rd</sup>** | 163/485 | 1.19 (0.92-1.54) | 1.20 (0.91-1.57) |
| **4<sup>th</sup> highest score** | 142/504 | 1 (reference) | 1 (reference) |
| DS |  |  |  |
| **1<sup>st</sup> lowest score** | 195/452 | 1.50 (1.17-1.93) | 1.53 (1.18-2.00) |
| **2<sup>nd</sup>** | 199/447 | 1.55 (1.21-1.99) | 1.53 (1.18-1.99) |
| **3<sup>rd</sup>** | 166/481 | 1.20 (0.93-1.55) | 1.25 (0.96-1.64) |
| **4<sup>th</sup> highest score** | 144/502 | 1 (reference) | 1 (reference) |

**Legend:** reference quartile (4<sup>th</sup> quartile) is the PGS category with the highest polygenic load for MDD/DS at the most significant threshold.
¥adjusted for age, sex, genotyping platform and 7-principal components. MDD: Major depressive disorder, DS: Depressive symptoms, PGS: polygenic score, OR: odds ratio

**Figure 2.**
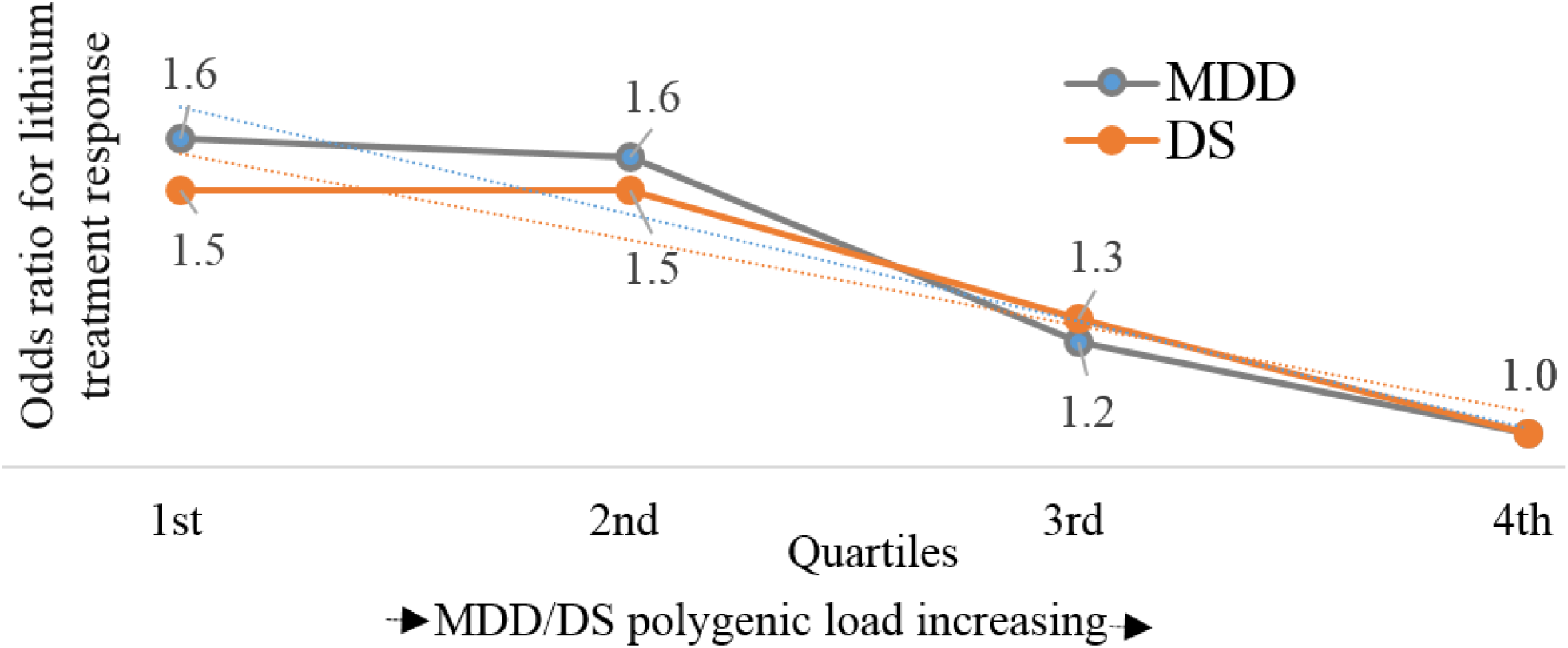
Odds ratios (ORs) for favorable treatment response to lithium for patients with BD in the low depression polygenic load quartiles (1^st^ to 3^r^), compared with patients in the highest depression polygenic load quartile (4^th^), estimated at the most significant p-value thresholds (nD = 02586).

Exploratory analyses were conducted to investigate whether the associations between MDD/DS PGSs and lithium response are driven by BD type 1 or type 2 (type 1 n=2,044; type 2 n=506). The observed inverse effects for MDD and DS PGS in the entire sample remained statistically significant for BD type 1 patients (eTable 3) only. For the BD type 2 group, an opposite non-significant trend for MDD PGSs, i.e. higher MDD loading was associated with *better* response to lithium, was observed (eTable 3).

### Cross-trait meta-analysis of GWAS on lithium treatment response and GWAS on MDD and depressive symptoms yields 7 significant loci

Subsequent to the PGS analysis, we performed a SNP-based cross-trait meta-analysis by combining the summary statistics for the GWASs on: a) MDD and lithium treatment response; and b) DS and lithium treatment response — with the aim of identifying individual genetic variants implicated in the genetic susceptibility to both depression traits and lithium treatment response. These analyses yielded 7 loci with p-values below the genome-wide significance level (p<5 × 10^−8^). These loci, and their nearby genes, were rs2327713: *PUM3* [OMIM: 609960], rs7134419: *KSR2* [OMIM: 610737], rs59659806: *RASGRP1* [OMIM: 603962], rs7405404: *ERCC4* [OMIM: 133520], rs11657502: *MYO18A* [OMIM: 610067], rs8099160: *DCC* [OMIM: 120470], rs6066909: *ARFGEF2* [OMIM: 605371] (Figure 3, Table 2)

**Table 2.**
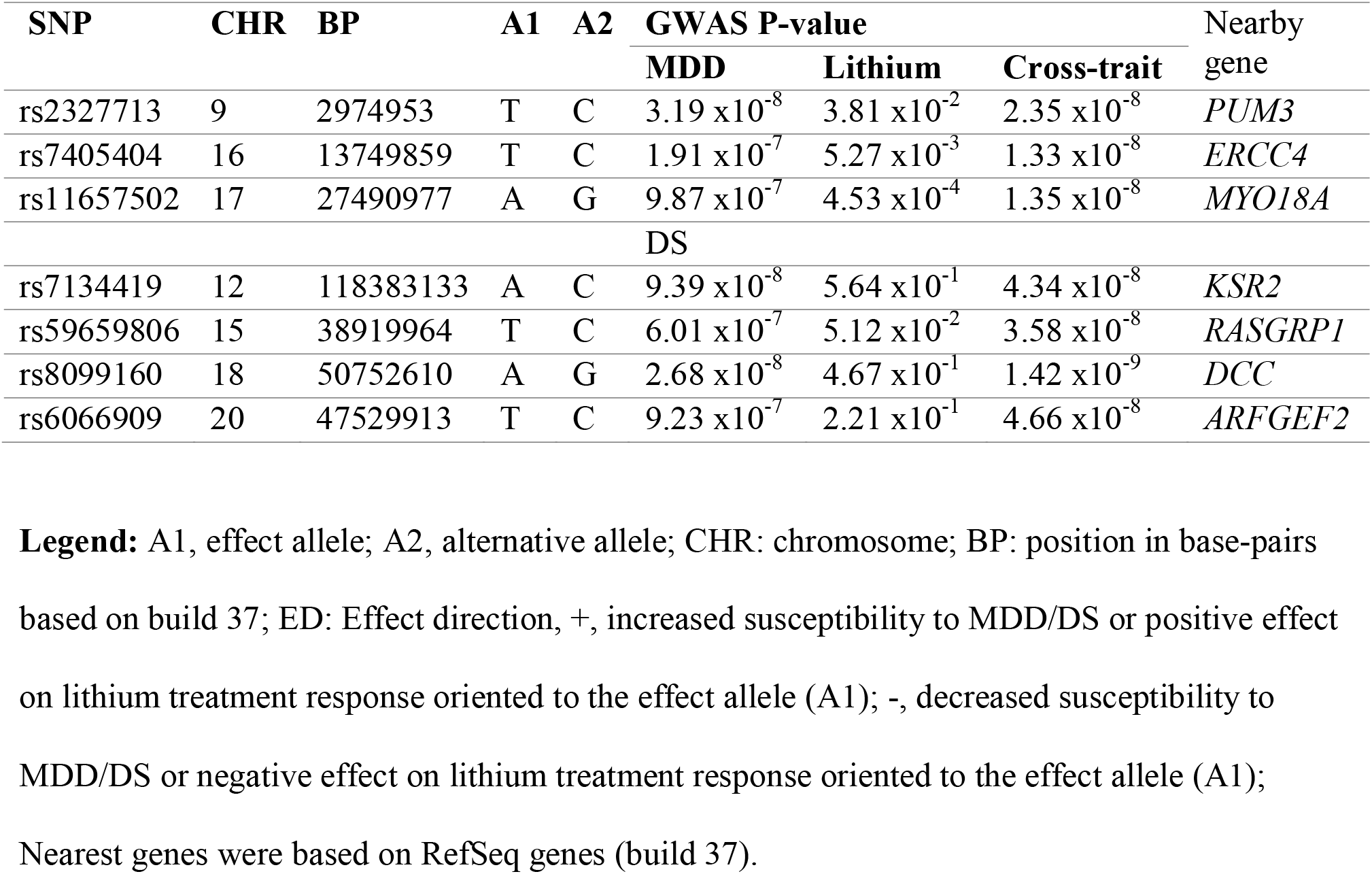
Loci resulting from a cross-trait meta-analysis of GWASs for lithium treatment response in patients with BD patients and GWAS for MDD and DS (cross-trait p<5 × 10^−8^).

| SNP | CHR | BP | A1 | A2 | GWAS P-value |  |  | Nearby gene |
| --- | --- | --- | --- | --- | --- | --- | --- | --- |
|  |  |  |  |  | MDD | Lithium | Cross-trait |  |
| rs2327713 | 9 | 2974953 | T | C | $3.19 \times 10^{-8}$ | $3.81 \times 10^{-2}$ | $2.35 \times 10^{-8}$ | *PUM3* |
| rs7405404 | 16 | 13749859 | T | C | $1.91 \times 10^{-7}$ | $5.27 \times 10^{-3}$ | $1.33 \times 10^{-8}$ | *ERCC4* |
| rs11657502 | 17 | 27490977 | A | G | $9.87 \times 10^{-7}$ | $4.53 \times 10^{-4}$ | $1.35 \times 10^{-8}$ | *MYO18A* |
| DS |  |  |  |  |  |  |  |  |
| rs7134419 | 12 | 118383133 | A | C | $9.39 \times 10^{-8}$ | $5.64 \times 10^{-1}$ | $4.34 \times 10^{-8}$ | *KSR2* |
| rs59659806 | 15 | 38919964 | T | C | $6.01 \times 10^{-7}$ | $5.12 \times 10^{-2}$ | $3.58 \times 10^{-8}$ | *RASGRP1* |
| rs8099160 | 18 | 50752610 | A | G | $2.68 \times 10^{-8}$ | $4.67 \times 10^{-1}$ | $1.42 \times 10^{-9}$ | *DCC* |
| rs6066909 | 20 | 47529913 | T | C | $9.23 \times 10^{-7}$ | $2.21 \times 10^{-1}$ | $4.66 \times 10^{-8}$ | *ARFGEF2* |

**Legend:** A1, effect allele; A2, alternative allele; CHR: chromosome; BP: position in base-pairs based on build 37; ED: Effect direction, +, increased susceptibility to MDD/DS or positive effect on lithium treatment response oriented to the effect allele (A1); -, decreased susceptibility to MDD/DS or negative effect on lithium treatment response oriented to the effect allele (A1); Nearest genes were based on RefSeq genes (build 37).

**Figure 3.**
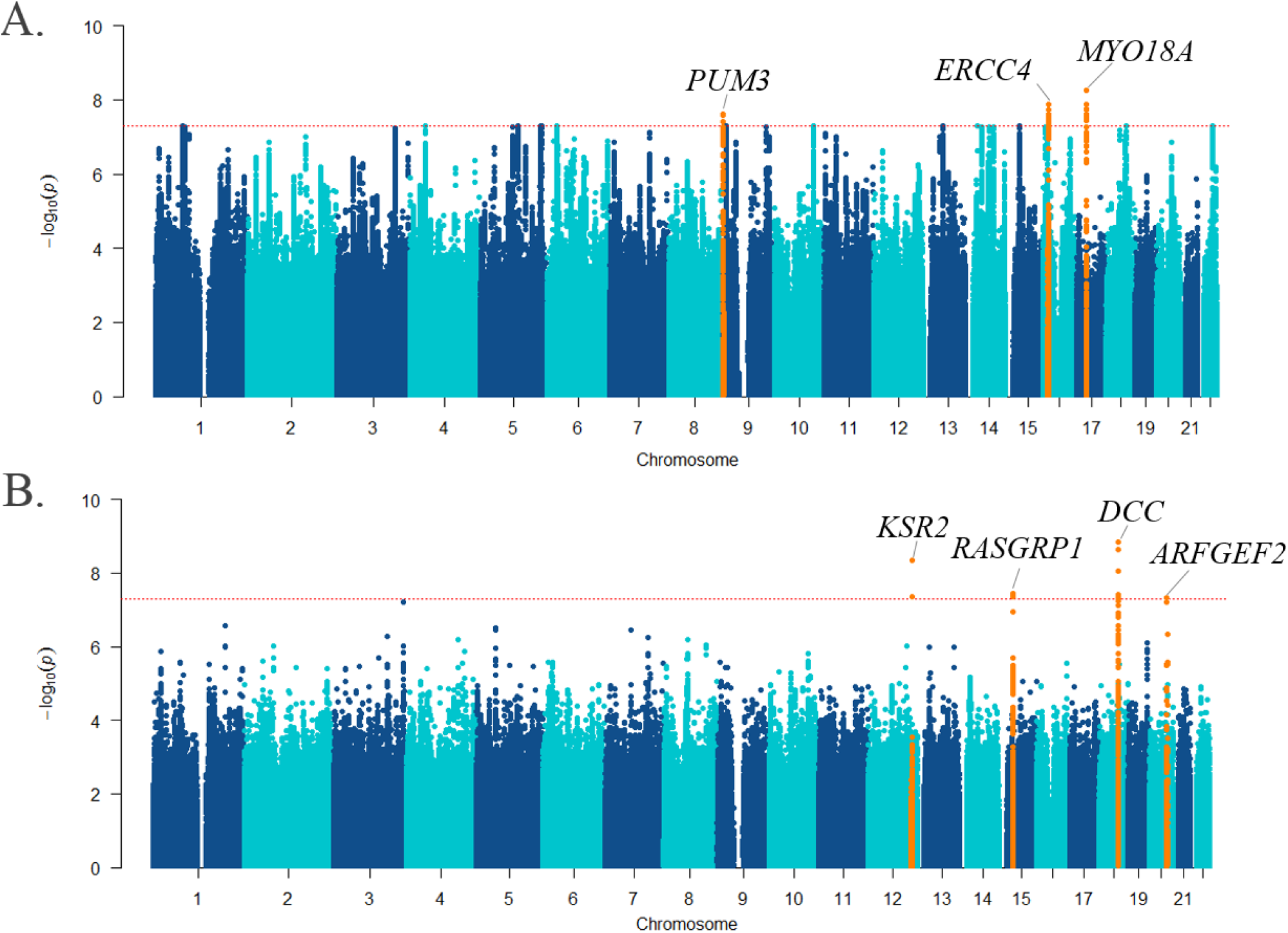
Manhattan plot of the cross-trait meta-analyses of GWASs on lithium treatment response and A) MDD; and B) DS. The loci that showed genome-wide significance are highlighted (orange), and their nearest genes indicated (top). The -log10 (cross-trait p-value) is plotted against the physical position of each SNP on each chromosome. The threshold for genome-wide significance (cross-trait p-value<5 × 10^−8^) is indicated by the red dotted horizontal line.

### Functional and biological characterization of genetic loci associated with lithium response and MDD/depressive symptoms

To characterize the functional implications of the SNPs identified by cross-trait meta-analysis, we first explored the functional genetic context of these variations by examination of SNPs with high linkage disequilibrium, characterization of their nearby hosting genes, and eQTL lookup from published databases. This approach yielded a list of 39 genes with potential functional significance [eTable 1]. Second, we investigated the biological roles of these 39 genes using IPA^®^ analysis.

**eTable 1.**
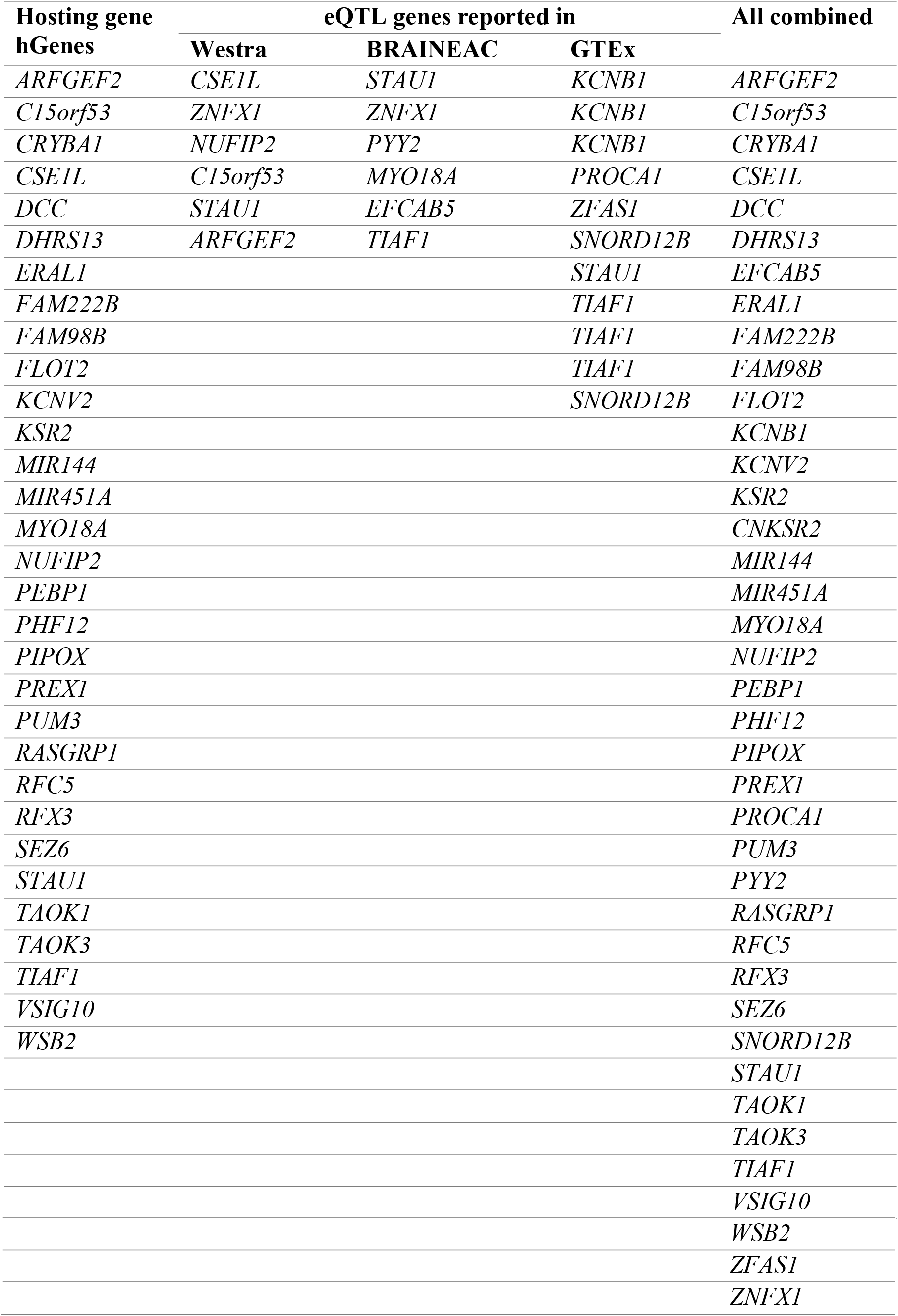
**Combined list of genes used for an input in the Ingenuity Pathway Analysis (IPA)**.

**eTable 2.** Molecules within the three-top significant functional networks

| Molecules in Network | Score | Focus Molecules |
| --- | --- | --- |
| **1, 2-dipalmitoylphosphatidylcholine, ACVR1C, CAMK2N2, CCNA1, CD300LD, Cxcl3, DCC, DEFB104A/DEFB104B, DSCAM, ERK, IL17RD, Insulin, Jnk, KCNB1, KCNB2, KCNV2, KSR2, L-lactic acid, Mapk kinase, MMP14, MUC8, N, N-dimethylarginine, PEBP1, PREX1, PROCA1, RASGRP1, RFX3, RFX6, S100A12, TAC4, TAOK3, TIAF1, TRIB3, VTCN1, ZFAS1** | 27 | 12 |
| **ARFGEF2, ARL5A, catalase, CDC42SE2, CREBBP, EFCAB5, ELAVL1, ETS2, FAM160B1, IMPA1, ITGB1, KLF7, KPNA2, MAP2K3, mir-144, mir-451, MSANTD3, NUFIP2, PIPOX, PUS7, SERTAD3, SLC25A44, SMAD4, SOX12, STK35, SUSD6, TAOK1, TBX4, TCR, TSPAN9, TSPAN31, TULP4, VSIG10, WSB2, ZNFX1** | 21 | 10 |
| **Alyref2, AVEN, CNKSR2, CSE1L, DHRS13, EED, EEFSEC, FAM98B, FLOT2, H2BFS, HIST1H2BN, HIST1H4E, HIST2H3D, HNRNPU, IMPA1, KIF11, KRBA2, MCM2, MYO18A, NXF1, OBSL1, PCID2, PHF12, PUM3, RFC5, RRP9, RRP15, STAU1, SURF6, TANC1, TUBA4A, TUBA4B, WDR43, ZBED8, ZNF142** | 21 | 10 |

**eTable 3.** The association of PGS for depression traits (MDD and DS) and lithium treatment response in patients with BD type 1 versus BD type 2 at different GWAS p-value thresholds (P_T_).

| **MDD PGS** |  | **BD type 1** |  | **BD type 2** |  |
| --- | --- | --- | --- | --- | --- |
| $P_T$ | No of SNPs | P-value | $R^2$ (%) | P-value | $R^2$ (%) |
| $P_T < 1 \times 10^{-4}$ | 852 | $1.83 \times 10^{-1}$ | 0.1% | 0.43 | 0.2% |
| $P_T < 1 \times 10^{-3}$ | 2987 | $1.09 \times 10^{-1}$ | 0.2% | 0.98 | 0.0% |
| $P_T < 1 \times 10^{-2}$ | 11892 | $4.34 \times 10^{-4}$ | 0.8% | 0.85 | 0.0% |
| $P_T < 5 \times 10^{-2}$ | 31712 | $2.94 \times 10^{-5}$ | 1.2% | 0.50 | 0.1% |
| $P_T < 0.1$ | 47716 | $8.86 \times 10^{-5}$ | 1.0% | 0.50 | 0.1% |
| $P_T < 0.2$ | 70784 | $1.08 \times 10^{-3}$ | 0.7% | 0.39 | 0.2% |
| $P_T < 0.3$ | 88552 | $2.61 \times 10^{-3}$ | 0.6% | 0.40 | 0.2% |
| $P_T < 0.4$ | 102503 | $1.33 \times 10^{-3}$ | 0.7% | 0.33 | 0.2% |
| $P_T < 0.5$ | 114071 | $1.72 \times 10^{-3}$ | 0.6% | 0.35 | 0.2% |
| $P_T < 1.0$ | 147596 | $2.97 \times 10^{-3}$ | 0.6% | 0.37 | 0.2% |
| **DS PGS** |  | **BD type 1** |  | **BD type 2** |  |
| $P_T$ | No of SNPs | P-value | $R^2$ (%) | P-value | $R^2$ (%) |
| $P_T < 1 \times 10^{-4}$ | 229 | $4.08 \times 10^{-2}$ | 0.3% | $2.67 \times 10^{-1}$ | 0.3% |
| $P_T < 1 \times 10^{-3}$ | 1352 | $1.18 \times 10^{-2}$ | 0.4% | $1.08 \times 10^{-1}$ | 0.7% |
| $P_T < 1 \times 10^{-2}$ | 8057 | $7.26 \times 10^{-3}$ | 0.5% | $6.26 \times 10^{-2}$ | 0.9% |
| $P_T < 5 \times 10^{-2}$ | 25918 | $5.51 \times 10^{-2}$ | 0.2% | $4.68 \times 10^{-2}$ | 1.0% |
| $P_T < 0.1$ | 41865 | $1.46 \times 10^{-1}$ | 0.1% | $4.86 \times 10^{-2}$ | 1.0% |
| $P_T < 0.2$ | 65634 | $1.54 \times 10^{-1}$ | 0.1% | $5.51 \times 10^{-2}$ | 0.9% |
| $P_T < 0.3$ | 83845 | $1.99 \times 10^{-1}$ | 0.1% | $5.48 \times 10^{-2}$ | 0.9% |
| $P_T < 0.4$ | 98217 | $1.61 \times 10^{-1}$ | 0.1% | $5.16 \times 10^{-2}$ | 1.0% |
| $P_T < 0.5$ | 109888 | $1.95 \times 10^{-1}$ | 0.1% | $4.96 \times 10^{-2}$ | 1.0% |
| $P_T < 1.0$ | 143345 | $2.16 \times 10^{-1}$ | 0.1% | $4.42 \times 10^{-2}$ | 1.0% |

IPA^®^ identified cellular development and cellular growth and proliferation as the top cellular functions associated with the 39 genes. These associations were driven by only a handful of genes, including micro RNA (miR) -144, miR-451, regulatory factor X3 (RFX), phosphatidylinsositol-3,4,5-trisphosphate dependent Rac exchange factor 1 (PREX1), and RAS guanyl releasing protein 1 (RASGRP1).

The top IPA^®^-identified functional networks containing dataset genes pointed to ‘hub’ functions for insulin, ERK, JNK (network 1), ELAV like RNA binding protein 1 (ELAV1)(network 2), and nuclear RNA export factor 1 (NXF1) (network 3), [eFigures 1 AC, eTable 2]. The IPA^®^ top hits for upstream regulators were potassium voltage-gated channel subfamily A member 1 (KCNA1) and leucine-2-alanine encephalin, both of which impact on the dataset gene potassium voltage-gated channel subfamily B member 1 (KCNB1) (Table 3).

**Table 3.** Top upstream regulators for genes identified in the cross-trait meta-analyses

| Upstream Regulator | Molecule type | P-value of overlap | Target molecules in dataset | Pathway for the regulator |
| --- | --- | --- | --- | --- |
| KCNA1 | ion channel | 9.31X10<sup>-4</sup> | KCNB1 | Voltage-gated potassium channel (function) |
| leucine-2-alanine enkephalin | chemical drug | 2.79X10<sup>-3</sup> | KCNB1 | opioid peptide (selective delta opioid receptor agonist) with analgesic properties |
| rubitecan | chemical drug | 3.72X10<sup>-3</sup> | PEBP1 | A semisynthetic agent with antitumor and antiviral properties |
| CAMKK2 | kinase | 7.43X10<sup>-3</sup> | KCNB1 | AMPK Signaling; Calcium Signaling; Dopamine-DARPP32 Feedback in cAMP Signaling |
| EGR2 | transcription regulator | 8.69X10<sup>-3</sup> | FLOT2, TAOK1 | Adipogenesis pathway |
| DAB1 | other | 8.35X10<sup>-3</sup> | DCC | Reelin Signaling in Neurons |

**Legend:** <sup>a</sup>P-values were adjusted by Benjamini & Hochberg (BH) method<sup>80</sup>. The p-value of overlap was determined at BH adjusted p-value $\leq 0.01$ .

**eFigure 1.**
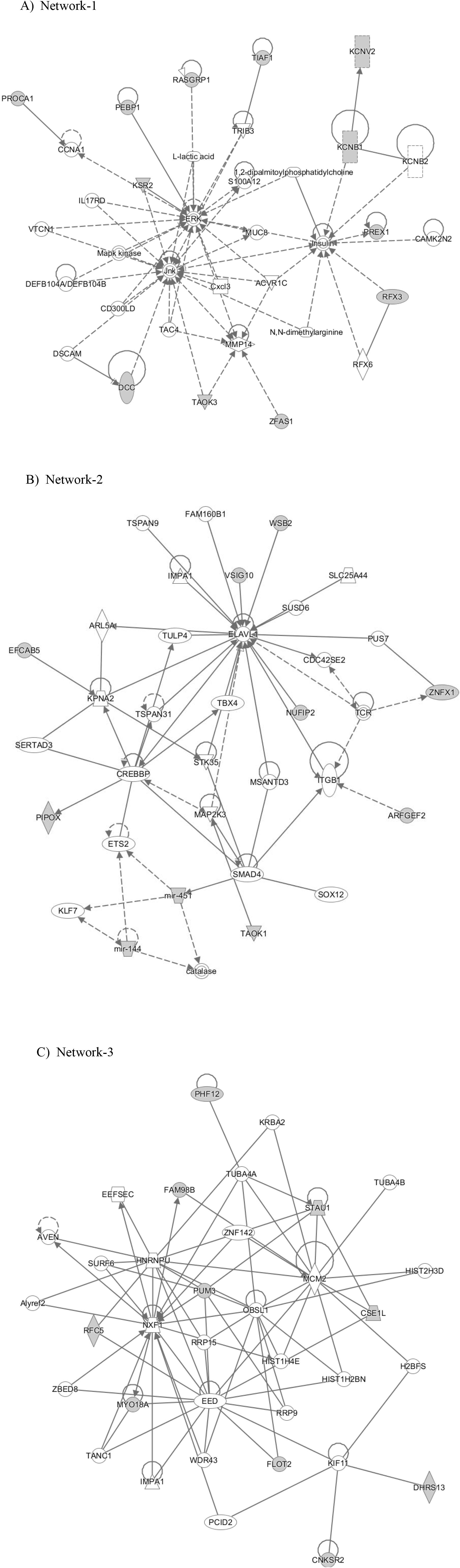
Top IPA^®^ functional network of genes associated with lithium response in BD and MDD/depressive symptoms. **eFigure 1 legend:** IPA generates the network using a proprietary algorithm, and included genes that could contribute to the network, even if they were not contained in the original dataset.

## Discussion

Our study represents the first direct evidence of an association between a genetic predisposition for depression and poorer response to lithium treatment in patients with BD. Using PGS analyses of genetic variants related to MDD and DS, we found that BD patients with low genetic loading for these variants were about 1.6 times *more* likely to have favorable long-term outcomes following lithium treatment compared to BD patients with high MDD/DS genetic loading. To explore which genes might functionally drive these effects, we carried out a cross-trait meta-analysis of lithium response and MDD/DS. Pathway analyses of variants associated with both traits implicated roles for voltage-gated potassium channels, for insulin-related mechanisms, for the ERK and JNK signaling pathways, and for the micro RNAs miR-144 and miR-451.

Our findings could form part of a genetic explanation for the previously described clinical observations in relation to mania, depression and lithium response in BD^6,7,27-34^ and supports the notion that lithium responsiveness could be associated with a ‘core’ bipolar phenotype in the *Kraepelinian* form of manic depression^34,44^. Such a concept is further supported by our previous finding of an inverse association of lithium response and schizophrenia PGS in BD^16^.

Although these results have to be interpreted with caution due to smaller subgroup sample sizes, the exploratory analyses indicate that BD type 2 patients may differ from type 1 patients with regards to the depression PGS on lithium response association. A non-significant trend for *improved* lithium treatment response was found in type 2 patients with high MDD PGS. Genetically, differences between type 1 and type 2 BD cohorts have been suggested ^45,46^, and type 2 patients show substantially higher genetic overlap with MDD^47^. Lithium’s effectiveness as an adjunct antidepressant treatment for people with treatment-resistant MDD is well established^48-54^; therefore, our finding raises the intriguing possibility of MDD-specific mechanisms of action, which might be different from the mechanisms underlying the more ‘anti-manic’ response in BD type 1.

Our cross-trait GWAS meta-analysis yielded 7 loci that exceeded a genome-wide significance level of 5×10^−8^. Amongst the nearby genes of these loci, the *DCC* gene and its encoded netrin 1 receptor has previously been shown to play an important role in mediating axonal growth in developing human brain^55,56^. Additionally, genetic variation within the *DCC* gene has previously been shown to be associated with depressive symptoms^36^.

Based on the 7 loci identified by cross-trait GWAS meta-analysis, we generated a list of 39 functionally related genes by examination of SNPs with high linkage disequilibrium, characterization of their nearby hosting genes, and eQTL lookup from published databases (eTable 1). Functional exploration of these genes by IPA^®^ implicated the voltage-gated potassium channel (K^+^v), KCNA1 as a top upstream regulator (Table 3). The family of K^+^v proteins plays a role in the regulation of the excitability of neurons, and genetic variations in these channels have been linked to epilepsy^57,58^. Recent experiments with inducible pluripotent stem cells from patients with BD suggest that neural hyperexcitability could be a core pathophysiological trait in BD, which is reversible by lithium in a subset of patients^59,60^. Remarkably, regulation of KCNA1 gene expression by lithium was shown to be involved in these ‘therapeutic’ lithium effects^59^. Therefore, a role for K^+^v’s in the genetic architecture of lithium response in BD appears plausible.

Further functional characterization by IPA^®^ suggested that genes regulating insulin homeostasis could be important mediators of the MDD-lithium relationship (eFigure 1). These genes included regulating factor X3 (RFX3), Phosphatidylinositol 3,4,5-trisphosphate-dependent Rac exchanger 1 (PREX 1), and K^+^v subfamily B member 1(KCNB1). Interestingly, previous clinical studies have shown that BD patients with impaired glucose tolerance or diabetes mellitus type 2 are over 8-times less likely to benefit from lithium and have an overall less favorable illness course ^19^.

Functionally, the genes *RFX3, PREX1*, and *KCNB1* are involved in insulin regulation in various ways. The transcription factor RFX3 is required for the differentiation and function of insulin-producing, mature pancreatic beta cells, and regulates the beta-cell promotor of the glucokinase gene^61^. Interestingly, *RFX3* variants were also implicated in a recent GWAS examining sleeplessness/insomnia^62^, a condition with aetiological relationships to BD^63^, and depression^64^. Variants of the PREX1 gene on chromosome 20q12-13.1were associated with increased risk of diabetes mellitus type 2 and increased BMI in a cohort of European Americans^65^, through mechanisms are yet insufficiently understood^66^. Variants of the KCNB1 gene in humans are associated with increases of waist to hip ratio, fasting insulin, and triglycerides, as well as decreased insulin sensitivity^67,68^. Mechanistically, the KCNB1-encoded Kv2.1 and other K^+^v are important for the fine-tuning of the release of cellular insulin and other hormones or neurotransmitters, and have both inhibitory (through re-polarization of the membrane potential^69^) and stimulating (through interaction with the soluble N-ethylmaleimide-sensitive factor attachment protein receptor (SNARE) complex)^70-73^ effects on exocytotic mechanisms. A previous study in rats suggested that treatment with lithium directly stimulates the expression of SNARE protein in brain tissue^74^.

Genetic variations in genes regulating ERK and JNK expression were identified as additional contributors to the effects of MDD PGS on lithium response. Belonging to a family of protein kinases in the mitogen-activated protein kinases (MAPK) pathway, these molecules are highly interactive with insulin-signaling mechanisms. Previous evidence has indicated that MAPK and insulin signaling could be activated by lithium, to enhance insulin-stimulated glucose transport and glycogen synthesis^75^. Additionally, lithium is known to stimulate MAPK-mediated neurite growth, neuronal survival, and neurogenesis^76^, and regulate circadian rhythms^77^. Therefore, it is possible that variation in MAPK-associated genes interferes with these potentially therapeutic effects of lithium.

MicroRNAs (miRNAs) regulate messenger RNA (mRNA) translation in a sequence-specific manner and are emerging as critical regulators of central nervous system plasticity. We found that genetic effects on miR-144 and miR-451 expression could play a role in mediating lithium response in BD. Previous animal studies have shown that lithium treatment in vivo induces changes in miRNA expression, specifically miR-144^78^. It is possible that variations of miRNA genes influence their contribution to lithium’s therapeutic mechanisms.

The main limitation of our study is that PGSs for MDD and DS explain only a small proportion of the variance in lithium treatment response (~1%), and have on their own no utility as clinical tests. Our cross-trait analysis provides a clue for a potential genetic overlap; however, no formal pleiotropy analyses were employed to confidently conclude about the effect of each genetic variant on the phenotypes tested. In addition, our pathway analysis findings are of an explorative nature and have not been validated on the transcript- or protein level or with experimental procedures in cellular models. Further, the current version of the Alda scale assesses only overall lithium efficacy but not effects specific to predominant illness polarity. Availability and incorporation of such information would have refined our results. The centrality of insulin-associated pathways in our findings could be a result of high representation of these genes within curated tools such as IPA^®^. However, these tools are powerful for hypothesis generation and indicate plausible molecular targets to be tested. Since our sample size already detected significant effects, it is likely that in the future, an increased sample size will further improve the predictive power of PGSs^79^.

In conclusion, we demonstrated that high genetic loadings for MDD and DS are predictive of unfavorable long-term response to lithium in patients with BD. Our study underscores the potential of PGS analysis to contribute to predictive models for medication response in psychiatry, and to uncover novel molecular pathways that drive these effects. While our findings, in isolation, are not yet ripe for clinical applications, they could serve as a component of multimodal predication models incorporating clinical and other biological data. The results of our study support clinical observations that have pointed to better lithium responsiveness in a BD subtype characterized by predominantly manic features. The study raises the possibility that mood-stabilizing- and anti-depressant properties of lithium are mediated through separate biological mechanisms.

## Acknowledgements

The authors are grateful to all patients who participated in the study and we appreciate the contributions of clinicians, scientists, research assistants and study staffs who have helped with patient recruitment, data collection and sample preparation for the studies. We are also indebted to the members of the ConLi^+^Gen Scientific Advisory Board (http://www.conligen.org/) for critical input over the course of the project.

The analysis of this study was carried out using the high-performance computational capabilities of the University of Adelaide, Phoenix supercomputer https://www.adelaide.edu.au/phoenix/.

